# Chronic corticosterone administration induces negative valence and impairs positive valence behaviors in mice

**DOI:** 10.1101/593632

**Authors:** Andrew Dieterich, Prachi Srivastava, Aitesam Sharif, Karina Stech, Benjamin A. Samuels

## Abstract

Behavioral approaches utilizing rodents to study mood disorders have focused primarily on negative valence behaviors associated with potential threat (anxiety). However, for disorders such as depression, positive valence behaviors that assess reward processing may be more translationally-valid and predictive of antidepressant treatment outcome. Chronic corticosterone (CORT) administration is a well-validated pharmacological stressor that increases negative valence behaviors (David et al., 2009; Gourley et al., 2008a,b; Gourley et al., 2012; Olausson et al., 2013). However, whether chronic stress paradigms such as CORT administration also lead to deficits in positive valence behaviors remains unclear. We treated male C57BL/6J mice with chronic CORT and assessed both negative and positive valence behaviors. We found that CORT induced negative valence behaviors associated with anxiety in the open field and NSF. Interestingly, CORT also impaired instrumental acquisition, reduced sensitivity to a devalued outcome, reduced breakpoint in progressive ratio, and impaired performance in probabilistic reversal learning. Taken together, these results demonstrate that chronic CORT administration at the same dosage both induces negative valence behaviors associated with anxiety and impairs positive valence behaviors associated with reward processing. These data suggest that CORT administration is a useful experimental system for preclinical approaches to studying stress-induced mood disorders.

**Significance Statement:** Chronic exposure to stress can precipitate mood disorders such as anxiety and depression. However, most studies focus on the effects of chronic stress on increasing negative affect behaviors. Elucidating how chronic stress impacts translationally-valid positive valence behaviors is less studied. Here, we show that chronic pharmacological stress induces negative affect behaviors associates with anxiety and impairs reward-related, positive valence behaviors in mice.

## Introduction

Mood disorders are a major burden on society and are extremely prevalent. An estimated 21.4% of adults in the United States experience a mood disorder at some point in their lives (Harvard Medical School National Comorbidity Survey), and approximately 16.2 million adults experience a major depressive episode each year (Substance Abuse and Mental Health Services Administration). Major depressive disorder is comprised of a gain of negative affect and a loss of positive affect. Specific symptoms can include depressed mood and/or anhedonia (Kessler et al., 2003). Anhedonia is the lack of feelings of pleasure and the positive experience from reward (Der-Avakian and Markou, 2012). Most preclinical rodent research focuses solely on assessing behaviors associated with the gain of negative affect. However, rodent behavioral approaches that assess anhedonia and/or loss of positive affect may prove to be more translationally relevant as relatively similar behaviors can be performed in rodents and humans (Der-Avakian et al., 2015). This conservation across species suggests that increased adoption of utilizing positive valence behaviors in rodents might significantly accelerate therapeutic development.

Chronic stress can precipitate mood disorders and several distinct chronic stress paradigms are widely used to study how stress impacts negative valence behaviors and neural circuitry in rodents. However, relatively few studies have assessed whether these chronic stress paradigms also impact positive valence behaviors. Chronic corticosterone (CORT) administration is one such paradigm that can mimic the effects of stress on negative valence behaviors (Zhao et al., 2008; David et al., 2009). CORT is the rodent stress hormone analog of cortisol in humans and is the major output of the hypothalamus-pituitary-adrenal (HPA) axis, which is hyperactive in humans with major depressive disorder and in rodents exposed to chronic stress (Gotlib et al., 2008; Boyle et al., 2005). Specifically, chronic CORT administration induces a negative valence phenotype, characterized by decreases in center measures in the open field and increased latency to eat in the novelty suppressed feeding test (David et al., 2009). Taken together, these negative valence behaviors that are more associated with potential threat/anxiety than depression suggest that chronic CORT administration can induce a negative affective state.

Here we show that chronic CORT administration not only induces negative valence behaviors, but also impairs positive valence behaviors. Following instrumental training, we utilized several positive valence behavioral tests, including outcome devaluation, progressive ratio, and probabilistic reversal learning, to assess the effects of chronic CORT on components of reward processing. Overall, CORT administration impaired instrumental acquisition, reduced sensitivity to a devalued outcome, reduced breakpoint in progressive ratio, and impaired performance in probabilistic reversal learning. Taken together, these data suggest that chronic stress paradigms, such as CORT administration, that are already validated with negative valence behaviors may also be useful for studying translationally relevant positive valence behaviors.

## Materials and Methods

### Mice and drug treatment

All experiments were conducted in compliance with NIH laboratory animal care guidelines and approved by Rutgers University Institutional Animal Care and Use Committee. Adult male C57BL/6 mice (age 7 weeks) (Jackson Labs, Bar Harbor, ME) were divided into Vehicle and Corticosterone (CORT) treatments. Vehicle-treated mice (n=20) received beta-cyclodextrin dissolved in their drinking water, while CORT-treated mice (n=30) received CORT (35 μg/mL) (Sigma-Aldrich, St. Louis, MO) and beta-cyclodextrin (4.5 mg/mL) (Sigma-Aldrich, St. Louis, MO) dissolved in their drinking water (5 mg/kg/day CORT). After 4 weeks of this treatment, mice were food-restricted and maintained at 85% −93% of their free-feeding body weight. To ensure the mice maintained this body weight they were weighed daily and given standard lab chow at least one hour after completing testing each day. Training conditions were collapsed into Vehicle (n=19) and CORT (n=26) treatment, as after initial instrumental acquisition there were no differences between training conditions on behavior. A total of 5 mice (n=1 Vehicle; n=4 CORT) died or did not reach criterion for acquisition, and were excluded from all further analyses. A smaller, separate cohort (n=10 for both Vehicle and CORT treatments) of adult male C57BL/6 mice was used for limited-training outcome devaluation and contingency degradation tests. In the negative valence cohort (n=20 for both Vehicle and CORT treatments), adult male C57BL/6 mice were tested in the open field (OF), novelty-suppressed feeding (NSF), and forced swim (FST) tests after 4 weeks of CORT or Vehicle (5mg/kg/day) administration.

### Open Field

Vehicle (n=20) and CORT (n=20) mice were placed in a corner of Plexiglas open field (OF) chambers measuring 43 × 43 cm, and Motor Monitor software (Kinder Scientific, Poway, CA) was used to measure distance traveled (cm), and both entries and time in the center of the OF, detected with infrared photobeams on the walls, and the center defined as a square 11 cm from each wall of the OF. Center behaviors were interpreted as measures of anxiety. This same cohort was used for additional negative valence behavior tests, including NSF and FST.

### Novelty-Suppressed Feeding

Vehicle (n=20) and CORT (n=20) mice were food-deprived 18 hours before novelty-suppressed feeding (NSF) testing. Mice were placed in a clean standby cage 30 minutes before NSF testing. Mice were placed in the corner of a novel, brightly-lit NSF chamber with a single food pellet placed in the center, for a 6-minute test. The NSF is a test assessing the conflict between motivation to eat while in a food-deprived state, and the aversive nature of the novel, brightly-lit arena. Latency to approach the pellet and take a bite was recorded. After the 6-minute test, mice were placed in their home cage for 5 additional minutes, and the weight and latency to eat a food pellet were recorded.

### Forced Swim

Vehicle (n=20) and CORT (n=20) mice were placed in forced swim (FST) chamber containing room temperature water for 6 minutes, and Motor Monitor software (Kinder Scientific, Loway, CA) was used to assess immobility, measured as 6 or less beam breaks in 5 seconds, during the final 4 minutes of the test.

### Operant Chambers

A large, separate cohort of Vehicle (n=20) and CORT (n=30), and a second, smaller cohort of Vehicle (n=10) and CORT (n=10) mice were trained and tested in standard mouse operant chambers (Med Associates, Fairfax, VT) housed in sound-attenuating cubicles, in a designated behavioral testing room. The operant chambers were coupled to a power control and interface connected to a computer running the Med-PC IV software (Med Associates, Fairfax, VT). The operant chambers contained two retractable response levers on one wall, and two 20 mg food pellet hoppers attached by Y-tubing to a single delivery port between the levers. Grain-based food pellet reinforcers (Bio-Serv, Flemington, NJ) were delivered from each hopper into the delivery port for the mouse to consume.

### Operant Conditioning

Mice were first habituated in two 30-minute sessions to the operant chambers, with only the house light on and fan running in the chamber. No levers were ejected into the chamber and no pellets were delivered into the food port. Then the mice were magazine-trained with pellets automatically delivered every 30 to 60 seconds. Following this, the levers were ejected into the chambers and the mice were trained to lever press in order to receive food pellet reinforcers, in addition to receiving the automatically-delivered pellets. One lever was ejected at a time, which alternated throughout each acquisition session as the mice learned to respond on both on a continuous reinforcement (CRF) schedule. After two days, no pellets were automatically dispensed, and the mice conditioned to respond on the CRF schedule until they reached criterion, which was 30 responses on both levers in consecutive sessions. Responses in each session and total sessions to reach criterion was recorded, to determine if CORT throughout training impairs instrumental acquisition.

### Extended-Training Outcome Devaluation

The larger cohort of mice (n=19 Vehicle; n=26 CORT) first completed extended-training, satiety-specific outcome devaluation testing to determine if chronic CORT affects responding for a devalued outcome. One hopper, counterbalanced across mice, was filled with chocolate-flavored (Bio-Serv) pellets, while the other continued to deliver standard grain-based food pellets. Thus, responding on one of the two levers led to delivery of a chocolate-flavored pellet, and responding on the other led to delivery of a grain-based pellet. Mice were subjected to two sessions of each of the following in succession: CRF, Random Ratio 5 (RR5) Schedule, RR10, and RR20. For each random ratio schedule, 5, 10, or 20 lever presses, on average, were required for reinforcer delivery, respectively. This protocol of extended-training has been used previously (Dias-Ferreira et al., 2009). After RR20 training, the mice progressed to satiety-specific devaluation of either the grain or chocolate pellets, as they were given free access to one of the pellet types in a fresh cage for one hour. Immediately after this they were given a 5-minute extinction test session where responses on both levers (associated with grain and chocolate pellets) were recorded but no reinforcers delivered. If the mouse received free access to the grain pellets, the lever associated with the grain pellets was considered the devalued condition, and the lever associated with the chocolate pellets was considered the valued condition. If the mouse received free access to the chocolate pellets, the lever associated with the chocolate pellets was the devalued condition, and the lever associated with the grain pellets was the valued condition. Lever presses on both sides were quantified and expressed as a ratio of responses per minute compared to the final day of RR20 training. This gave normalized response rates for both valued and devalued conditions.

### Extended-Training Contingency Degradation

Mice were then re-trained on a RR20 schedule on both levers until responding stabilized, still for both grain-based and chocolate-flavored pellets, counterbalanced by side. They then continued on one active lever on the RR20 schedule, while the second lever became inactivate, as pellets were delivered based on the rate of responding on the active lever, to degrade responding for that outcome. When responding on the degraded lever had robustly decreased in both Vehicle and CORT mice, a 5-minute extinction test session was conducted with both levers ejected into the chamber, and responses on both levers were recorded, to determine if chronic CORT impacts responding for a degraded contingency.

### Limited-Training Outcome Devaluation

As over-training causes a shift to habitual control of responding, in the limited-training outcome devaluation test a separate cohort of Vehicle (n=10) and CORT (n=10) mice was briefly habituated to and trained on a CRF schedule to lever press on one response lever for the grain-based food pellets. Mice were then trained on a VR2 schedule of reinforcement for three days, where 1, 2, or 3 responses led to reinforcer delivery, after which they were given free access to either the reinforcer pellets (devalued condition), or standard lab chow (valued condition) for one hour prior to a 5-minute extinction test where no reinforcers were delivered but lever presses were recorded. Mice completed both conditions in separate sessions, with two days in between testing to test if CORT affects sensitivity to a devalued outcome after limited instrumental training.

### Limited Training Contingency Degradation

For contingency degradation, Vehicle (n=10) and CORT (n=10) mice from the limited-training outcome devaluation experiment were re-trained on a VR2 schedule but now on both response levers, for three days, or until responding stabilized, as described previously (Gourley et al., 2012). Then, one of the levers was retracted and responding on the other lever (non-degraded condition) was recorded in one VR2 session, counterbalanced across mice. In the next session the other lever was ejected into the chamber (degraded condition) but was not paired with the reinforcer. Rather, pellets were dispensed at the same rate as in the previous day’s VR2 session, to degrade responding on the degraded lever. A 10-minute extinction test session was conducted the following day to assess if CORT reduces sensitivity to a degraded contingency.

### Progressive Ratio

After completing extended-training contingency degradation, the cohort of Vehicle (n=19) and CORT (n=26) mice was then trained on a VR3 schedule for three days, where 2, 3, or 4 lever presses were required for reinforcement. After this the mice completed three separate progressive ratio test sessions, to test for motivation to expend effort. In each, the requirement to obtain a food pellet reinforcer increased linearly (1, 5, 9, X+4) until the mice ceased to respond for 5 minutes, or until 4 hours. The highest ratio completed, or breakpoint, was recorded, which provides a measure of motivation to work to obtain a reinforcer. Total active lever presses were also measured in each session. After completing all three progressive ratio tests, the large cohort progressed to reversal learning.

### Reversal Learning

Vehicle (n=19) and CORT (n=26) mice were then trained in 30-minute sessions on a reversal learning procedure, where they were reinforced on a CRF schedule for each response made on either the left or right lever (correct), counterbalanced between mice, while the other lever (incorrect) was not reinforced. After 8 consecutive responses, recorded as a completed reversal on the correct lever, the correct and incorrect levers switched, and the previously incorrect, non-reinforced lever was now the correct, reinforced lever. With an incorrect response, the counter started over and the mouse was required to make an additional 8 consecutive correct response to complete the reversal and switch the contingencies. Correct and incorrect levers switched with each completed reversal in each reversal learning session. Correct responses, omissions (no response made within 10 seconds), incorrect responses, and completed reversals were measured. Following five sessions of reversal learning, the mice completed three different probabilistic reversal learning (PRL) test sessions, which were each separated by two days of reversal learning sessions.

### Probabilistic Reversal Learning

PRL was conducted as previously described (Ineichen et al., 2012; Bari et al., 2010) in Vehicle (n=19) and CORT (n=26) mice. The procedure for PRL is similar to reversal learning, except PRL can be divided in to three separate sessions involving different reinforcement probabilities. In each, the correct lever is reinforced on only 70%, 80%, or 90% of correct presses, while the incorrect lever is reinforced on 10%, 20%, or 30% of the incorrect presses, respectively, which is referred to as pPCR, or the probability of punishment for a correct response. Therefore, the three PRL sessions are referred to as 0.1, 0.2, and 0.3 pPCR. These probabilities of reinforcement induce misleading positive and negative feedback. A lose-shift occurred when a mouse switches to respond on the incorrect lever immediately after making a correct, but not reinforced response. A win-stay occurs when a mouse continues to respond on the correct lever after a correct response made in the immediately preceding trial. These are measures of negative and positive feedback, respectively, and were assessed along with completed reversals for each of the three PRL test sessions.

### Experimental Design and Statistical Analysis

The effect of chronic CORT treatment on positive valence behaviors was assessed using separate one-way ANOVAs, t-tests, or Kaplan-Meier Survival analyses. Separate cohorts of Vehicle and CORT-treated mice were used for separate experiments. Vehicle (n=20) and CORT (n=2-0) mice were used in the negative valence cohort. Vehicle (n=10) and CORT (n=10) mice were used in the limited-training outcome devaluation cohort. Vehicle (n=19) and CORT (n=26) mice were used for the extended training outcome devaluation, progressive ratio, and probabilistic reversal learning tests. Planned multiple comparisons were made to determine significance between groups.

## Results

Exposure of rodents to chronic stress induces negative valence behaviors associated with potential threat/anxiety. Therefore, to confirm that chronic CORT administration (5mg/kg/day) mimics the effects of chronic stress, we assessed behavior in the open field (OF), novelty suppressed feeding (NSF), and forced swim test (FST) in a cohort of C57BL/6J male mice (n=20 Vehicle; n=20 CORT) (Figure 1a). Similar to previous reports (David et al., 2009), unpaired t-tests revealed that chronic CORT administration reduces percent time spent in the center (*t*(37) = 3.5, *p* < 0.001, Figure 1d), and entries into the center of the OF (*t*(37) = 5.5, *p* < 0.001, Figure 1c), as well as overall locomotor activity in the OF (*t*(37) = 4.5, *p* < 0.001, Figure 1b). Furthermore, chronic CORT increased latency to eat in NSF (Kaplan-Meier survival, X^2^(2, N = 36) = 16, *p* < 0.001, Figure 1e). However, similar to previous reports (David et al 2009), chronic CORT did not affect immobility in antidepressant-sensitive FST (*t*(37) = 0.017), *p* = 0.987, Figure 1f). Taken together, these results confirm that chronic CORT administration induces negative valence behaviors associated with potential threat/anxiety in the OF and NSF.

**Figure 1.**
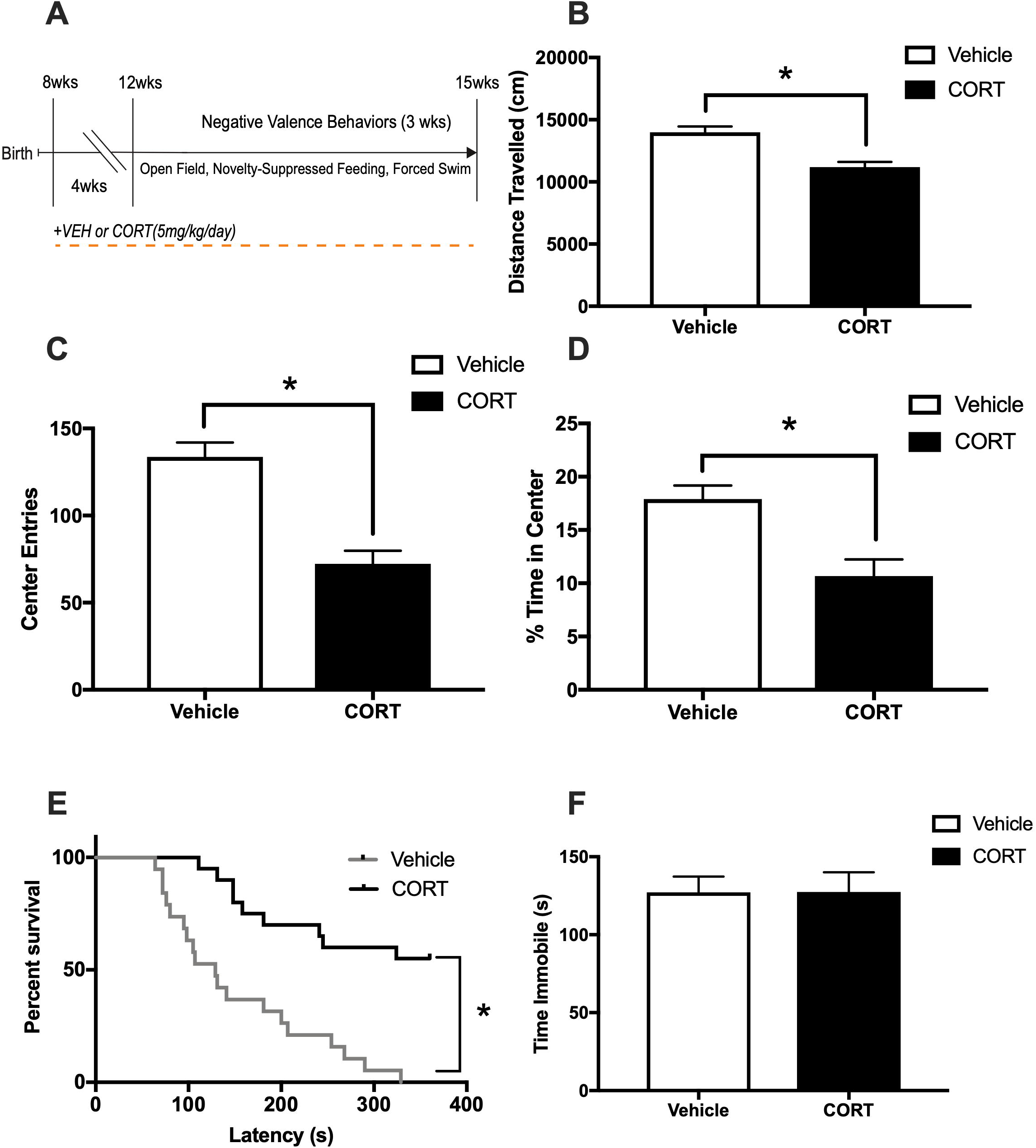
Chronic CORT increases negative valence. (A) Vehicle (n=20) or CORT (n=20) C57BL/6 mice were administered CORT or Vehicle (5mg/kg/day) in their drinking water for 4 weeks prior to negative valence behaviors, including the open field (OF), novelty suppressed feeding (NSF), and forced swim (FST) tests. (B) In the OF, CORT reduced locomotor activity, measured as distance traveled (cm) in a 30-minute session (*t*(37) = 4.5, *p* < 0.001). (C) CORT also reduced center entries in the OF, (*t*(37) = 5.5, *p* < 0.001), (D) as well as percent time in the center (*t*(37) = 3.5, *p* < 0.001), both measures of anxiety behavior in the OF. (E) In the NSF assay, chronic CORT increased latency to take a bite of a food pellet in the center of the brightly-lit arena under a food-deprived state (Kaplan-Meier survival, *X*^2^(2, N = 36) = 16, *p* < 0.001). (F) In the FST, CORT had no effect on immobility, indicative of behavioral despair (*t*(37) = 0.017), *p* = 0.987), a commonly-used depressive test to assess antidepressant response. Values plotted are mean ± SEM. *p < 0.001. Taken together, chronic CORT induces a robust negative valence behavioral phenotype characterized by increased anxiety in the OF and NSF, while not increasing immobility in the FST.

While the effects of chronic stress paradigms on negative valence behaviors are widely studied in rodents, much less is known as to whether chronic stress impacts positive valence behaviors. Therefore, we first assessed the effects of CORT on instrumental acquisition using a CRF schedule with 30 lever presses in two consecutive sessions considered to be acquisition (Figure 2a). Using a distinct cohort of C57BL/6J mice than the negative valence behaviors, we tried two different protocols, where CORT was either administered throughout training (n=10 Vehicle, n=15 CORT) or after the completion of training (n=10 Vehicle, n=15 CORT). A two-way ANOVA revealed a main effect of training on sessions to acquisition (*F*(1, 36) = 4.88, *p* = 0.034) and an interaction between training and CORT administration (*F*(1,36) = 8.726, *p* = 0.006) (Figure 2b). Multiple comparisons revealed that CORT administration throughout training resulted in significantly more sessions to acquisition than mice that were administered CORT after the completion of training (*p* = 0.003) or were administered vehicle throughout training (*p* = 0.035). A two-way repeated measures ANOVA revealed a main effect of day on responses during training (*F*(3, 126) = 28.36, *p* < 0.001), a main effect of CORT administration, (*F*(3, 42) = 8.50, *p* < 0.001), and a significant day x CORT administration interaction, (*F*(9, 126) = 2.30, *p* = 0.02) (Figure 2c). Mice that received CORT throughout training made significantly less responses than mice that received CORT after the completion of training on all 4 training days (Day 1: *p* = 0.023, Days 2, 3, and 4: *p* < 0.001) and mice that received vehicle throughout training on Days 3 (*p* < 0.001) and 4 (*p* = 0.029). Taken together, these data demonstrate that CORT administration throughout training impairs instrumental acquisition compared to Vehicle and to CORT administration after training.

**Figure 2.**
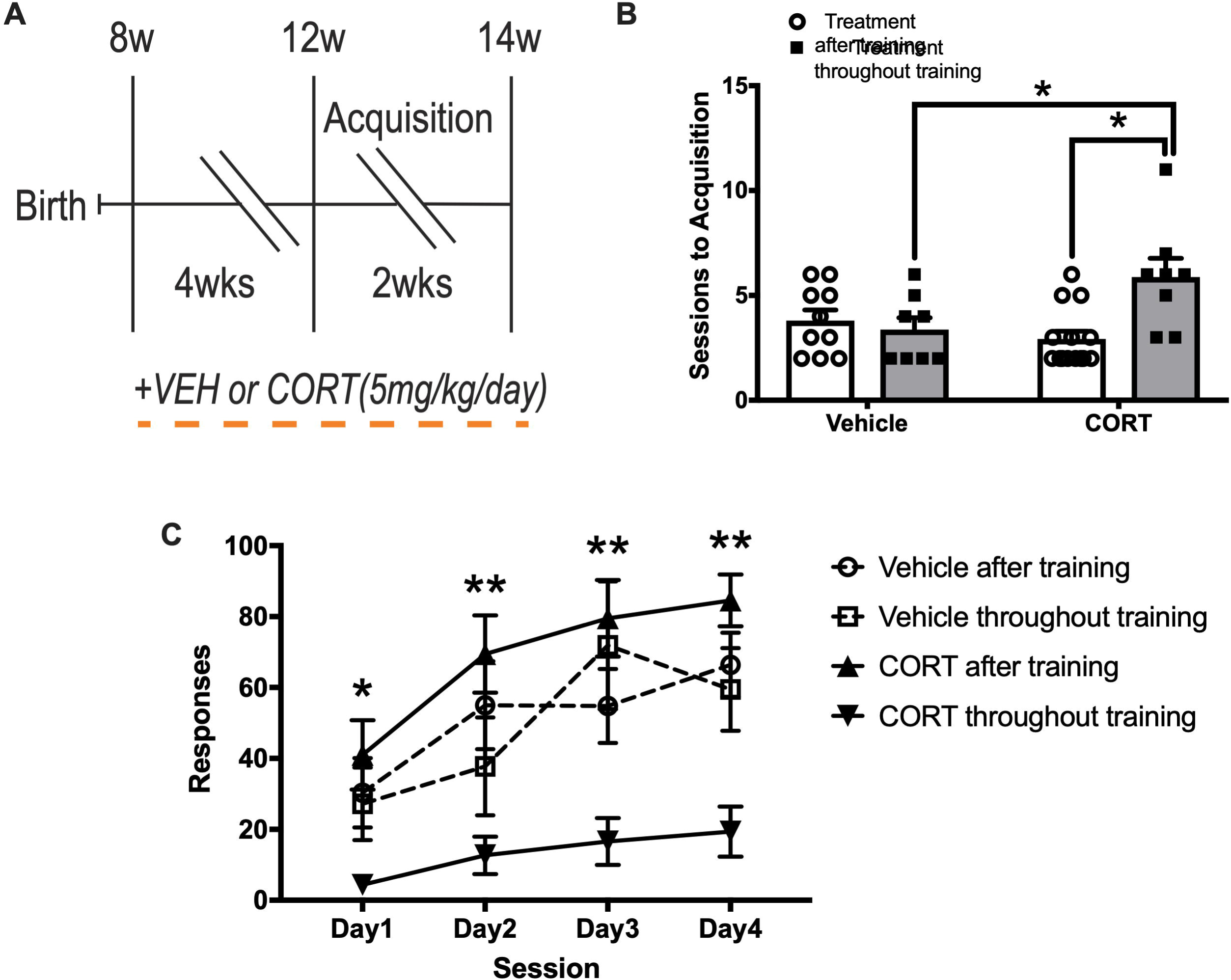
Chronic CORT impairs instrumental acquisition. (A) Vehicle (n=10, both training conditions) and CORT (n=15, both training conditions) mice were administered CORT or Vehicle (5mg/kg/day) in their drinking water for 4 weeks prior to instrumental acquisition (Vehicle or CORT throughout training), or after instrumental acquisition (Vehicle or CORT after training). (B) Sessions to acquisition (criterion defined as consecutive sessions with >= 30 lever presses on both response levers) was assessed in Vehicle and CORT mice throughout or after treatment. CORT throughout training mice required more acquisition sessions to reach criterion compared to CORT after training (*p* = 0.003), and Vehicle throughout training (*p* = 0.035). (C) Responses were recorded throughout acquisition and were reduced by CORT treatment throughout training compared to CORT treatment after training (Day 1: *p* = 0.023, Days 2, 3, and 4: *p* < 0.001) and compared to Vehicle treatment throughout training on Days 3 (*p* < 0.001) and 4 (*p* = 0.029). Values plotted are mean ± SEM. *p < 0.05, **p < 0.001. Taken together, CORT throughout training inhibits instrumental acquisition, reducing lever presses during acquisition, and increasing sessions to reach criterion for acquisition.

Since all mice reached the acquisition criterion, we collapsed Vehicle (n=19) and CORT (n=26) mice for the remainder of behavioral testing. We next assessed whether continued CORT administration affected extended-training satiety-specific outcome devaluation in mice that completed instrumental acquisition (Figure 3a). Briefly, mice went through an extended training protocol to press one lever for a grain-based food pellet and a distinct lever for a chocolate-flavored pellet. Following the completion of this extended training, mice were given free access to one of the pellet types in a new cage for one hour. Immediately after the free access they were given a 5-minute extinction test where responses on both levers were recorded but no reinforcers delivered. The pellet type they had free access to was considered devalued. For satiety-specific outcome devaluation, a two-way ANOVA for normalized (compared to the final RR20 session) lever pressing revealed no main effect of stress (*F*(1, 86) = 0.45, *p* = 0.504), value condition (*F*(1, 86) = 0.83, *p* = 0.365), or interaction (*F*(1, 86) = 0.67, *p* = 0.416) (Figure 3b). Vehicle (*p* = 0.518) and CORT (*p* > 0.999) mice similarly responded on valued and devalued levers during the extinction test. This indicates that CORT does not impact extended-training outcome devaluation. Mice were next tested in extended-training contingency degradation, to determine if chronic CORT affects responding on a degraded contingency. Mice were re-trained on a RR20 schedule on both levers, and then one lever was degraded by automatically delivering pellets at the rate of reinforcement from the other, non-degraded lever. After three days, mice lever-pressed in an extinction session to determine if CORT administration reduces sensitivity to a degraded outcome. For contingency degradation, a two-way ANOVA for normalized lever pressing revealed a main effect of degradation condition (*F*(1, 86) = 34, *p* < 0.001), but no effect of stress, (*F*(1, 86) = 0.018, *p* = 0.672), and no interaction (*F*(1, 86) = 0.92, *p* = 0.339 (Figure 3c)). Planned multiple comparisons indicate both Vehicle and CORT mice reduce responses on a degraded lever compared to the non-degraded lever (*p* < 0.001), suggesting this protocol produced a robust degradation effect in both treatments. Taken together, Vehicle and CORT mice both reduced responding for a reinforcer devalued by contingency degradation, suggesting that like outcome devaluation, CORT does not affect responding when compared to Vehicle.

**Figure 3.**
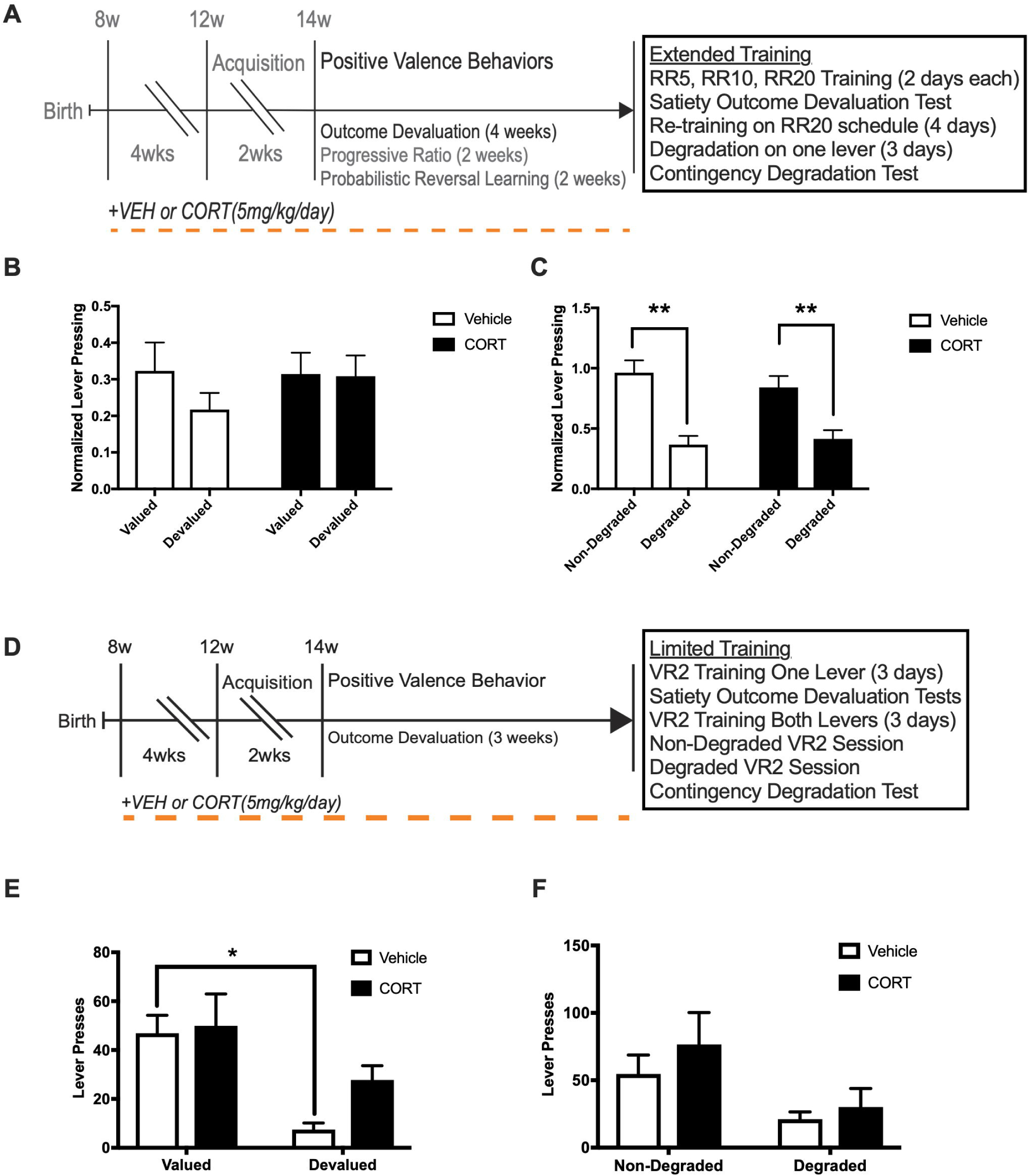
Chronic CORT impairs outcome devaluation after limited instrumental conditioning, but not extended training. (A) Training groups (treatment after and treatment throughout training) were collapsed into treatments (Vehicle and CORT) after initial instrumental acquisition. Mice were tested for satiety-specific outcome devaluation and contingency degradation after both extended and limited-training, to test for the effect of chronic CORT on sensitivity to a devalued or degraded outcome, and whether goal-directed or habitual mechanisms are guiding behavior. (B) Vehicle (n=19) and CORT (n=26) mice did not differ in normalized lever pressing in extinction test sessions after satiety-specific outcome devaluation (*p* = 0.262). (C) Vehicle (n=19) and CORT (n=26) mice similarly reduced lever pressing on a degraded response lever (*p* < 0.001 for both treatments), but not on a non-degraded lever (*p* > 0.05), suggesting extended-training similarly degraded responding on one lever in both treatments, as the lever was degraded for 3 degradation sessions. (D) Next, a separate cohort of Vehicle (n=10) and CORT (n=10) mice were trained in a shorter protocol before satiety-specific outcome devaluation and contingency degradation, to examine if the extended-training protocol led to over-training. (E) After 1-hour free access to reinforcer pellets (devalued), or standard lab chow (valued), limited-training led to a reduction in lever pressing in the devalued condition in Vehicle mice (*p* = 0.003) but not in CORT mice (*p* = 0.128), suggesting CORT reduces sensitivity to a devalued outcome after limited training. (F) However, limited training did not lead to reduced responding for a degraded outcome like shown with extended-training in either Vehicle (*p* = 0.163), or CORT (*p* = 0.091) mice. Values plotted are mean ± SEM. *p < 0.05, **p < 0.001. Taken together, chronic CORT reduces sensitivity to a devalued outcome after limited trained, without influencing contingency degradation, while extended-training shifts behavior to habitual responding in both treatments.

Since overtraining may have caused a shift to habitual control of responding in the extended-training condition, we also tried a limited-training protocol in a distinct cohort of C57BL/6J male mice (Figure 3d). Vehicle (n = 10), and CORT (n = 10) mice were trained on a CRF schedule to criterion (30 lever presses in a single session) on one lever and then completed three sessions on a VR2 schedule. Mice were tested in a 5-minute extinction session under two conditions and on separate days: after 30 minutes of free access to lab chow (valued), or free access to reinforcer pellets (valued). A two-way ANOVA revealed a main effect of value condition on responding during the extinction sessions (*F*(1, 32) = 15, *p* < 0.001), but a non-significant main effect of CORT administration (*F*(1, 32) = 2, *p* = 0.162), or interaction (*F*(1, 32) = 1.1, *p* = 0.295) (Figure 3e). Planned multiple comparisons indicated that the Vehicle mice show a significant devaluation effect (*p* < 0.003), while CORT mice did not show this effect (*p* = 0.128). Thus, chronic CORT administration impairs outcome devaluation, as CORT mice are insensitive to the devalued outcome. These mice were next assessed in a limited training contingency degradation test. A two-way ANOVA for the effect of contingency degradation on responding in an extinction session revealed a main effect of degradation condition (*F*(1, 30) = 7.6, *p* = 0.01), but no effect of CORT administration (*F*(1, 30 = 1.1, *p* = 0.296), or interaction (*F*(1, 30) = 0.2, *p* = 0.659) (Figure 3f). Planned multiple comparisons indicated no degradation effect for Vehicle (*p* = 0.163), or CORT (*p* = 0.091). These data demonstrate that chronic CORT reduces sensitivity to a devalued outcome when mice are briefly trained, while not affecting a degraded contingency between response and reinforcer.

We next assessed the effect of CORT administration on progressive ratio in the cohort previously exposed to the extended training to test if chronic CORT impairs motivation to expend effort for a reinforcer. To this end, Vehicle (n=19), and CORT (n=26) mice were trained on a VR3 schedule for three days, followed by three separate progressive ratio sessions where the response requirement to obtain a reinforcer increased linearly (1, 5, 9, X+4) for either 4 hours or until the mice ceased to respond for 5 minutes (Figure 4a). A two-way repeated measures ANOVA revealed a main effect of CORT administration on active lever presses (*F*(1, 43) = 65.77, *p* < 0.001), and a main effect of day on active lever presses (*F*(2, 86) = 8.08, *p* < 0.001), but a non-significant interaction (*F*(2, 86) = 0.96, *p* = 0.387) (Figure 4b). Bonferroni’s multiple comparisons test indicates CORT reduced active lever presses compared to Vehicle in all three progressive ratio sessions (*p* < 0.001). For the three progressive ratio sessions, a two-way repeated measures ANOVA revealed a main effect of CORT administration on last ratio reached (*F*(1, 43) = 56.80, *p* < 0.001), and a main effect of day on last ratio reached (*F*(2, 86) = 4.98, *p* = 0.009), but a non-significant interaction (*F*(2, 86) = 0.41, *p* = 0.668) (Figure 4c). Bonferroni’s multiple comparisons test indicates CORT reduced last ratio reached compared to Vehicle in all three progressive ratio sessions (*p* < 0.001). Therefore, chronic CORT treatment reduces motivation to lever press on a progressively increasing requirement, as CORT decreased breakpoint and total active lever presses.

**Figure 4.**
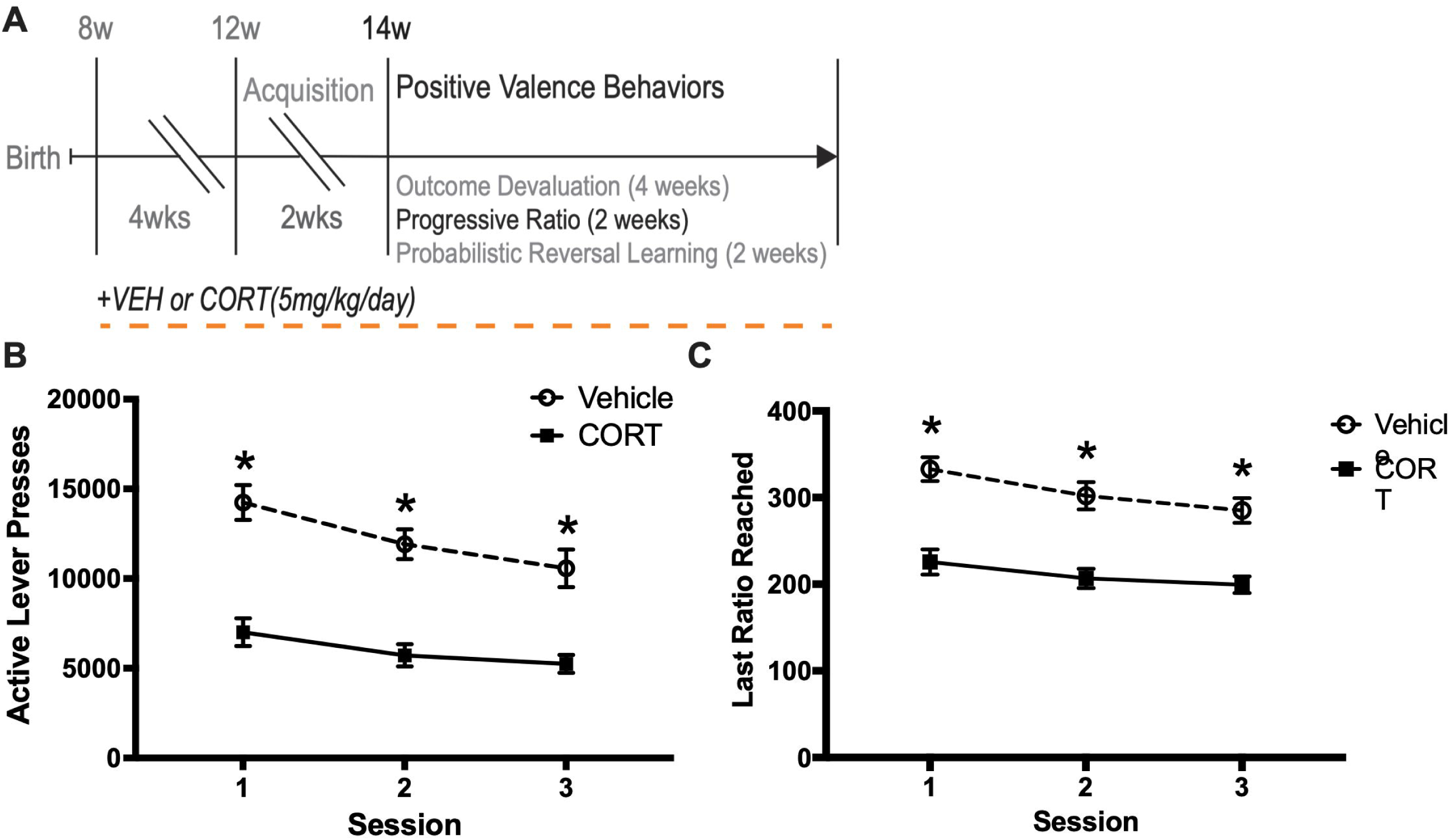
Chronic CORT reduces active lever presses and breakpoint ratio in the progressive ratio test. (A) After completing outcome devaluation and contingency degradation tests, Vehicle (n=19) and CORT (n=26) mice were trained for 3 sessions on a VR3 schedule and then tested in progressive ratio in 3 sessions separated by 2 days, under a linearly increasing, X+4 (1, 5, 9, X+4) progressive ratio schedule. (B) Chronic CORT reduced total active lever presses compared to Vehicle in all 3 sessions (*p* < 0.001). (C) Chronic CORT similarly reduced last ratio reached in all 3 sessions (*p* < 0.001). Values plotted are mean ± SEM. *p<0.001. Taken together, chronic CORT reduces motivation to expend effort for food pellet reinforcers by reducing lever presses and breakpoint in 3 progressive ratio tests.

Lastly, we assessed the effects of CORT administration on probabilistic reversal learning in the cohort previously exposed to the extended training outcome devaluation and the progressive ratio. Vehicle (n=19), and CORT (n=26) mice were trained for five days on a reversal learning protocol, where one lever was “correct” and reinforced, and the other lever was “incorrect” and not reinforced (Figure 5a). After 8 consecutive correct responses, the two levers switched. A two-way repeated measures ANOVA for completed reversals during the five days of reversal learning training revealed a main effect of CORT administration (*F*(1, 43) = 39.2, *p* < 0.001), a main effect of training day (*F*(4, 172) = 69.76, *p* < 0.001), and a significant interaction (*F*(4, 172) = 10.97, *p* < 0.001) (Figure 5b). Bonferroni’s multiple comparisons test indicated that CORT reduced completed reversals compared to Vehicle on training days 3, 4, and 5 (*p* < 0.001). Across the three sessions of probabilistic reversal learning, there were main effects of CORT administration (*F*(1, 43) = 26.37, *p* < 0.001), and probability of punishment for a correct response (pPCR) (*F*(2, 86) = 13.14, *p* < 0.001), on completed reversals, and a non-significant interaction *(F*(2, 86) = 0.97, *p* < 0.38 (Figure 5c). Planned Bonferroni’s multiple comparisons indicated CORT reduced completed reversals when pPCR was 0.1 (*p* < 0.001), and 0.2 (*p* = 0.01), but not when pPCR was 0.3 (*p* = 0.054). Across the three days of probabilistic reversal learning, there was a main effect of pPCR on the probability of making a Win-Stay (*F*(2, 86) = 67.43, *p* < 0.001), and a main effect of CORT administration (*F*(1, 43) = 9.054, p = 0.004), but the interaction between factors was non-significant (*F*(2, 86) = 0.27, *p* = 0.77) (Figure 5d). Planned Bonferroni’s multiple comparisons indicated CORT trended to reduce the probability of making a Win-Stay when pPCR was 0.1 (*p* = 0.052). Across the three days of probabilistic reversal learning, there were no main effects of pPCR (*F*(2, 86) = 1.147, *p* = 0.322) or treatment *F*(1, 43) = 0.1, p = 0.754) on probability of making a lose-shift (Figure 5e). These data demonstrate that CORT impairs cognitive flexibility in both reversal learning and probabilistic reversal learning tasks, as CORT mice failed to complete as many reversals as Vehicle mice during training, and in all three PRL sessions. Further, CORT may impact positive feedback sensitivity, as CORT trended to reduce win-stays when pPCR was 0.1.

**Figure 5.**
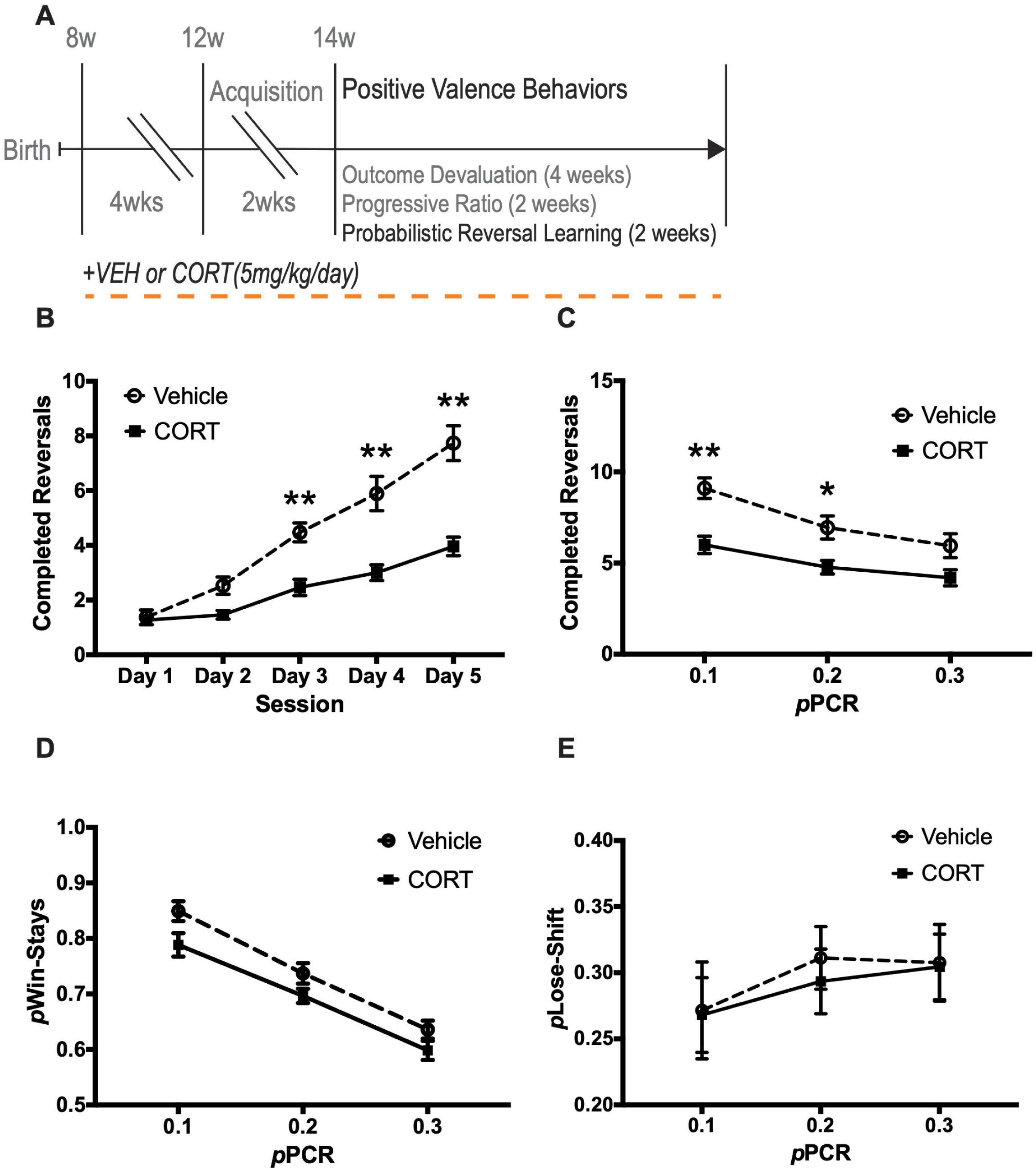
Chronic CORT impairs completed reversals in reversal learning and probabilistic reversal learning. (A) Vehicle (n=19), and CORT (n=26) mice were trained for five days on reversal learning, followed by 3 separate probabilistic reversal learning (PRL) test sessions, where the probability of punishment for a correct response (pPCR), was 0.1, 0.2, and 0.3, counterbalanced between mice. (B) Chronic CORT reduced completed reversals compared to Vehicle on reversal learning training days 3, 4, and 5 (*p* < 0.001). (C) Chronic CORT also reduced completed reversals in the 0.1 (*p* < 0.001) and 0.2 (*p* = 0.01) pPCR PRL test sessions, but not in the 0.3 pPCR PRL session (*p* = 0.38). (D) Chronic CORT trended to reduce the probability of making a win-stay in the 0.1 pPCR PRL session. (E) CORT has no effect on the probability of making a lose-shift in any of the PRL sessions. Values plotted are mean ± SEM. *p<0.05, **p<0.001. Taken together, chronic CORT impairs cognitive flexibility in both reversal learning and PRL sessions by reducing competed reversals. While non-significant, CORT also trends to reduce the probability of making a win-stay in the 0.1 pPCR PRL session (*p* 0.052).

## Discussion

Similar to previous reports (David et al., 2009), chronic CORT induced a robust negative valence behavioral phenotype in both open field and novelty suppressed feeding. However, here we show that chronic CORT administration also had several effects on positive valence behaviors including causing impairments in instrumental acquisition and limited-training satiety-specific outcome devaluation. CORT administration also reduced motivation to expend effort for reinforcer pellets in progressive ratio and reduced completed reversals in reversal learning and in PRL test sessions. Taken together, these data suggest that chronic CORT administration induces both an increase in negative affect and a decrease in positive affect. Our chronic CORT administration paradigm utilizes the lowest dose (5 mg/kg/day) possible that consistently induces negative valence behaviors (David et al., 2009). Our data here suggests that this same administration paradigm is also useful for studying the effects of chronic stress on translationally relevant positive valence behaviors.

Acquisition involves the formation of an action-outcome (A-O) contingency, as lever pressing leads to reward presentation. Devaluing this response via outcome devaluation or contingency degradation tests if behavior is controlled by a goal-directed (A-O) or stimulus-response (S-R), habitual mechanism. While non-stressed mice reduce responding for devalued food pellets (via contingency degradation), chronic CORT treated mice maintained their level of responding, indicating an S-R, habitual mechanism underlying this pattern of responding. Thus, chronic CORT reduces sensitivity to a devalued outcome in satiety-specific outcome devaluation. Chronic social defeat stress also reduces A-O decision-making, therefore reducing responding for a devalued reinforcer using a satiety-specific outcome devaluation (Dias-Ferreira et al., 2009). Therefore, also given our data here, it is possible that chronic stress paradigms such as CORT administration and social defeat stress originally designed for and validated using negative valence behaviors can also be used to determine the effects of chronic stress on positive valence behaviors.

Other studies have utilized subchronic and/or higher doses of CORT to assess effects on positive valence behaviors in rats and mice and have reported mixed results. One study found that subchronic CORT (at a dose ∼60% higher than we used) impairs instrumental acquisition (Olausson et al., 2013) while another found that subchronic CORT (at a dose similar to what we used) had no effect on instrumental acquisition (Gourley et al., 2012). Both of these studies utilized nose poke apertures for mice during the acquisition. Our chronic CORT administration protocol, which uses a dose well-validated for negative valence behaviors (5mg/kg/day) (David et al 2009) resulted in impaired instrumental acquisition using lever presses during the acquisition. Furthermore, subchronic CORT (at a dose similar to what we used) impaired both lithium chloride-paired outcome devaluation and also contingency degradation in mice utilizing nose poke apertures (Gourley et al., 2012). We found that chronic CORT impaired satiety-specific outcome devaluation in mice utilizing lever presses. However, we did not see a significant effect of CORT administration on contingency degradation. Extended-training mice showed significant effects of degradation, as lever pressing was reduced in the extinction test session, regardless of vehicle or chronic CORT treatment, and there was no effect of degradation in the limited-training cohort, regardless of vehicle or chronic CORT treatment. Finally, subchronic CORT (using doses either 40% higher or 540% higher than we used) reduced nose poke responding in a progressive ratio task (Gourley et al., 2008a,b). Importantly, to our knowledge the data presented here are also the first to assess the effects of CORT administration on probabilistic reversal learning.

Probabilistic reversal learning assesses behavioral flexibility, and the sensitivity to both positive and negative feedback (Ineichen et al., 2012; Bari et al., 2010). In a reversal learning procedure rodents respond to a correct lever for a reward pellet on 8 consecutive trials (Rychlik et al., 2017). Then, the rewarded lever switches sides, and the rodent must respond on the other lever to receive a reward. This contingency switches throughout the reversal learning session, and the number of completed reversals is considered a measure of cognitive flexibility. In the probabilistic versions of this test correct responses are not reinforced, while incorrect responses are reinforced at a determined probability. This provides positive and negative feedback, which are assessed with win-stays and lose-shifts during the task. We found that chronic CORT reduced completed reversals in both training and during probabilistic versions of the test, and that win-stays trended to be reduced in CORT mice when pPCR was 0.1. This novel finding further demonstrates that the CORT paradigm produces a robust blunting of positive valence throughout several behavioral tests.

Correspondence between human and rodent measures of reward learning, such as in the PRL task, is especially high (Der-Avakian et al., 2016). The human versions of PRL (Taylor Tavares et al., 2008; Cools et al., 2002) do not rely on self-reporting and the rodent versions of PRL were actually developed based on the parameters used in humans (Bari et al., 2010; Ineichen et al., 2012). In the human version of the PRL task subjects must learn to alter responding to a reversed contingency and ignore both positive and negative misleading feedback to maximize the probability of reward (Evers et al., 2005; Cools et al., 2002). Humans must visually discriminate between stimuli on a computer screen, which can be abstract patterns or lines of similar length (Evers et al., 2005). While they are rewarded for correct choices, the stimuli reverse just like in the rodent version of this task. Negative feedback occurs similarly as the rodent version, with the same pPCR of 0.1, 0.2, and 0.3. In agreement with our chronic CORT findings, depressed patients show greater focus on negative feedback in the PRL test (Taylor Tavares et al., 2008). While previous rodent work has assessed the effects of various antidepressant treatments on PRL (Bari et al., 2010; Ineichen et al., 2012; Drozd et al., 2019; Rychlik et al., 2016), this study provides the first evidence that a chronic stressor such as CORT leads to similar effects in PRL as to what is observed in depressed patients.

Mood disorders are mediated by a complex neural circuitry involving interconnected regions including the nucleus accumbens, medial prefrontal cortex, amygdala, ventral hippocampus, and ventral tegmental area (Russo and Nestler, 2013). Dysfunction in distinct circuits likely mediate the different symptoms observed in mood disorders. While significant effort has helped decipher the neural circuitry underlying negative valence behaviors, especially those associated with anxiety (Calhoon and Tye, 2015), less is known about the circuitry mediating positive valence behaviors. Therefore, future studies will need to also focus on deciphering the neural circuitry of positive valence and the effects of chronic stress on this circuitry.

## Acknowledgements

The authors would like to thank Dr. Mimi Phan and Mark Presker for assistance in setting up the operant chambers, and Dr. Mark West for helpful discussion. This work was funded by NIMH R01 MH112861 (BAS).

## Specific Contributions

A.D., P.S., A.S., and K.S. performed research. A.D. and B.A.S. designed research, analyzed data, and wrote the paper.

## Specific Contributions

**Extended Data Figure 2-1.** Vehicle (n=10) and CORT (n=15) treated mice were trained to acquire the instrumental response throughout treatment (5 mg/kg/day) or after treatment. Then the mice completed outcome devaluation, progressive ratio, and probabilistic reversal learning. For progressive ratio, a two-way ANOVA shows a main effect of treatment, *F*(3, 41) = 21.27, *p* < 0.001, session, *F*(2, 82) = 8.23, *p* < 0.001, but no interaction, *F*(6, 82) = 0.86, *p* = 0.531 for total active lever presses. Planned multiple comparisons indicate no differences between Vehicle (*p* = 0.746) or CORT (*p* = 0.951), indicating there were no difference between training groups. Values plotted are mean ± SEM. Thus, we collapsed them into respective treatments (n=19 Vehicle; n=26 CORT) for positive valence behavioral testing.

